# Microbubble-Mediated Cavitation Promotes Apoptosis and Suppresses Invasion in AsPC-1 Cells

**DOI:** 10.1101/2020.05.18.102152

**Authors:** J. Cao, H. Zhou, F. Qiu, J. Chen, F. Diao, P. Huang

## Abstract

This study aimed to identify the potential and mechanisms of microbubble-mediated cavitation in promoting apoptosis and suppressing invasion in cancer cells. AsPC-1 cells were used and divided into four groups: control group, microbubble-only (MB) group, ultrasound-only (US) group, and ultrasound combined with microbubbles (US+MB) group. Pulse ultrasound was used with a frequency of 360 kHz and an intensity of 0.7 W/cm^2^ for 1 min (duty rate=50%). Then cells in four groups were cultured for 24 h. Cell counting kit‑8 illustrated that US+MB could decrease cell viability. Western blot confirmed that US+MB increased cleaved Caspase‑3, Bcl-2-associated X protein (BAX), and decreased the B‑cell lymphoma‑2 (Bcl-2) levels. Besides, US+MB increased intracellular calcium ions and down-regulated cleaved Caspase-8. For proliferation ability, cells in US+MB group had a lower expression of Ki67 and the weakened colony formation ability. The transwell invasion assay showed that US+MB could decrease invasion ability in AsPC-1 cells. Further evidence showed that cells conducted with US+MB had the lower level of hypoxia-inducible factor-1α (HIF-1α) and Vimentin and the higher expression of E-cadherin than the other three groups. Finally, cells conducted with US+MB had less invadopodia formation. In conclusion, these results suggested that microbubble-mediated cavitation promoted apoptosis and suppressed invasion in AsPC-1 cells.

**Significance:** Cancer cells have a high rate of cell division and a high rate of cell division can speed up the development of hypoxia which lead to promote cancer invasion and metastasis. Here we used microbubble-mediated cavitation to generate a mechanical shock wave which could effectively kill cancer cells through physical damage and intrinsic signaling pathway. Furthermore, effective killing of cancer cells could restrain the development of hypoxia and prevent invasion ability of cancer cells.

## Introduction

Pancreatic cancer is the second leading cause of death from cancer in the digestive system, and the 5-year survival of patients with pancreatic cancer is <5% (1). The high mortality rate of patients with pancreatic cancer is mainly because of the insidious onset, invasive fast-growing, early metastasis, and therapy resistance. Cancer cells have a high rate of cell division and the mutated genes accelerate cell growth, leading to the loss of the normal cell cycle (2). Conventional treatments for pancreatic cancer include surgery, chemotherapy, and radiation therapy, aiming to kill tumor cells directly or induce cell apoptosis to reduce the number of tumor cells (3). Furthermore, a high rate of cell division also speeds up the development of hypoxia, which is considered as an important factor to induce epithelial-mesenchymal transition (EMT) and promote cancer metastasis (4, 5). EMT is the process that cells lose the characteristic of epithelial cells and obtain the potential of mesenchymal cells, which confers cancer cells the abilities of metastasis and therapy resistance (6). Hence, restraining the development of hypoxia and EMT is a key part in preventing cancer invasion.

Recently, the wider usage of ultrasound technology has led a deeper awareness of biological effect induced by ultrasound cavitation. Cavitation is the process that bubbles form and collapse with the change of local pressure, which can be divided into stable cavitation and inertial cavitation; the former induces a reversible, increased permeability of the cell membrane, while the latter generates a large amount of energy leading to the irreversible disruption of the cell membrane (7, 8). For the low gas solubility in the medium at room temperature, cavitation can be enhanced with the existence of microbubbles (9). Several researchers have reported the effect that the microbubble-mediated cavitation can induce apoptosis in cancer cells and be applied to cancer therapy in animals and at a cellular level. In animals, ultrasound combined with microbubbles can effectively restrain tumor growth (10, 11). At the cellular level, the potential of cavitation in promoting cell apoptosis has been examined in some cell lines (12–14). Researchers also have explored the mechanism underlying the cavitation-induced cell apoptosis, showing that increased levels of intracellular calcium and reactive oxygen species can eventually activate Caspase-3 and cause cell apoptosis (15–17). However, Caspase-3 can be activated through several ways, including the mitochondria-mediated pathway and death receptor-mediated pathway, and its upstream signals contain not only the increased intracellular calcium ions but also Caspase-8 (18). Therefore, cavitation-induced cell death remains to further explore mechanisms. Besides, there are very few reports confirming the ability of ultrasound coupled with microbubbles to inhibit cancer cell invasion. This study aimed to investigate the potential and mechanisms of ultrasound combined with microbubbles in promoting apoptosis and suppressing invasion in AsPC-1 cells.

## Materials and methods

### Cell culture, ultrasound equipment, and microbubbles

AsPC-1 cells were purchased from the Cell Bank of the Chinese Academy of Sciences (Shanghai, China), cultured in an RPMI 1640 medium (HyClone, USA) with 10% fetal bovine serum (Biological Industries, USA), and maintained in a humidified incubator at 37°C and 5% CO_2_. The cells were divided into four groups: control group, microbubble-only (MB) group, ultrasound-only (US) group, and ultrasound combined with microbubbles (US+MB) group. The SonoVue™ echo-contrast agent (Bracco, Italy) was used as microbubbles to enhance the effect of cavitation. This echo-contrast agent powder was suspended by 5 ml sterile saline and shaken well before using, about 2 × 10^8^ microbubbles per milliliter. The diameter of microbubbles in the suspension was between 2-8 μm. Add the microbubble suspension to MB group and US+MB group with the liquid volume fraction of 10% while add the same amount of sterile saline to control group and US group. The ultrasound treatment comprised a signal generator and amplifier (Model AG1010 Amplifier/Generator, T&C Power Conversion Inc., USA), a 360-kHz transducer (No. 715 Institute of China Shipbuilding Industry Group Co. Ltd., China), and a water bath. The water bath was used to reduce acoustic attenuation. The cell culture plate was placed vertically 5 cm over the transducer. The center acoustic pressure was 145 kPa measured by a hydrophone, and the center acoustic intensity was 0.7 W/cm^2^ calculated by the following equation (19):

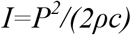

where *I* is the acoustic intensity in W/m^2^, *P* is the acoustic pressure in Pa, *ρ* is the medium density in kg/m^3^ (here we considered the density of the RPMI 1640 medium as equal to that of water, i.e., 1000 kg/m^3^), and *c* is the acoustic velocity in m/s (here we considered the acoustic velocity as equal to that in the water at room temperature, i.e., 1500 m/s). Cells in US group and US+MB group were exposed to the pulse ultrasound wave for 1 min with 60 cycles (duty rate=50%) while cells in control group and MB group were treated without ultrasound.

### Cell viability measurement

The cells were seeded at a density of 6000 cells per well in a 96-well cell culture plate and exposed to the different experimental conditions after adhering. The Cell Counting Kit-8 (CCK8) (Biosharp, China) was used to measure cell viability 24 h after the treatment. A 10 μl amount of reagent was added to each well, and the cells were incubated for 2 h before testing the absorbance at 450 nm. Each group set 8 wells to decrease the difference within each group. After obtaining the optical density (OD) values in four groups, the minimum and maximum values in each group were deleted and the rest OD values in each group were normalized by the average OD value in the control group.

### Colony formation assay

Colony formation reflects the characters of proliferation ability and population dependence. The cells were seeded at a density of 3000 cells per well in a 6-well cell culture plate. After adhering, the cells were exposed to the different experimental conditions (control, MB, US and US+MB) and cultured for 14 days. Crystal violet was used to stain cells and then phosphate buffered saline (PBS) to wash the crystal violet for three times. The ratio of violet area was calculated by ImageJ (National Institutes of Health, Bethesda, MD, USA) and the ratio was considered as the ability of cell colony formation.

### Western blot analysis

The cells were seeded at a density of 500,000 cells per well in a 6-well cell culture plate. After adhering, the cells were exposed to the different experimental conditions (control, MB, US and US+MB) and then cultured for 24 h. The cells were lysed using a RIPA lysis buffer, and the protein concentration was measured using a Bicinchoninic Acid Protein Quantification kit (Beyotime, China). The equal amount of protein from each group was loaded to the lane and separated by sodium dodecyl sulfate-polyacrylamide gel electrophoresis (SDS-PAGE). The protein was then transferred onto nitrocellulose membranes. Nonfat milk was used to block the transferred membrane for 1 h, after which the membranes were incubated with primary antibodies overnight at 4°C. Primary antibodies were listed below: Ki67 (Abcam 16667), β-actin (CW 0096M), β-tubulin (CW 0098M), B-cell lymphoma-2 (Bcl-2, CST 3498), Bcl-2-associated X protein (Bax, Abcam 263897), cleaved Caspase-3 (Abcam 13585), cleaved Caspase-8 (CST 9429), E-cadherin (CST 3195), hypoxia-inducible factor-1α (HIF-1α, CST 3716). On the next day, the membranes were washed with a mixture of tris-buffered saline and Tween 20 (TBST) and incubated with corresponding secondary antibodies for 1 h at room temperature. All membranes were exposed to a ChemiDoc MP Imager. Bands of Ki67, cleaved Caspase-3, Bcl-2, Bax, HIF-1α and E-cadherin were normalized to β-actin and bands of cleaved Caspase-8 were normalized to β-tubulin. Protein levels were quantified by ImageJ.

### Scanning electron microscopy

To observe morphological change of cells under different experimental conditions, scanning electron microscopy (SEM) was performed immediately after the cells were exposed to the different experimental conditions. Cell samples of the four groups were fixed with 2.5% glutaraldehyde solution and stored at 4°C for >4 h. They were then rinsed with phosphate buffered saline (PBS) and dehydrated with increasing concentrations of ethanol. After the dehydrated cells were dried, they were analyzed under a field-emission SEM.

### Cell apoptosis assay

The cells were seeded at a density of 500,000 cells per well in a 6-well cell culture plate. After adhering, the cells were exposed to the different experimental conditions (control, MB, US and US+MB) and then cultured for 24 h. Then, the cells were harvested with 0.25% trypsin enzyme without ethylene diamine tetraacetic acid (EDTA) and centrifugated for 5 min at the speed of 2000 r. p. m at room temperature. Then the cells were suspended with cold PBS and centrifugated for 5 min at the speed of 2000 r. p. m at room temperature. Then the cells were stained using an Annexin V-FITC/PI apoptosis detection kit (BD Pharmingen). After the cells were collected, 300 μl binding buffer was added to suspend cells and 5 μl Annexin V-FITC was added to the cell suspension with sufficient mixing. The cell suspension was stored away from light for 15 min at room temperature and 5 μl PI staining solution was added 5 min before testing. Annexin V (+)/PI (−) and Annexin V (+)/PI (+) cells were identified as cells in the early and late apoptosis stages, respectively. Annexin V (−)/PI (+) cells were considered as cell debris. The results of flow cytometry were analyzed using the Kaluza Analysis Software (Beckman Coulter Inc., CA, USA). Cells in Annexin V (+)/PI (−), Annexin V (+)/PI (+) and Annexin V (−)/PI (+) were considered as the apoptotic cells.

### Immunofluorescence analysis

The cell slides were collected 24 h after the experiment. Then, the cells were fixed with 4% polyoxymethylene and permeabilized with 0.2% Triton-X in PBS for 20 min. After blocking with 5% bovine serum albumin, the cells were incubated with primary antibodies at 4°C overnight. Primary antibodies were listed below: Ki67 (Abcam 16667), E-cadherin (CST 3195) and Vimentin (Abcam 16700). Then secondary antibodies conjugated with a fluorescence dye were applied for 1 h at room temperature. 4’,6-diamidino-2-phenylindole (DAPI) was also applied for nucleus staining. Then the slides were exposed to a fluorescent microscope. The ratio of the target protein area to the nucleus area was calculated by ImageJ. F-actin staining and β-tubulin staining were used to track morphological changes of the cytoskeleton. The cell slides were collected 24 h after the experiment. Then, the cells were fixed with 4% polyoxymethylene and permeabilized with 0.2% Triton-X in PBS for 20 min. After blocking with 5% bovine serum albumin, the cells were incubated with β-tubulin (CST 2146) at 4°C overnight and incubated with the corresponding secondary antibody conjugated with a fluorescence dye (Thermo A-21428) and phalloidin conjugated with a fluorescence dye (CST 8878) for 1 h at room temperature before the slides were exposed to a fluorescent microscope.

### Intracellular calcium measurement

To measure the intracellular calcium level, Fluo-8 AM (AAT Bioquest, USA) was used to bind with calcium ions inside the cells. The cell slides were collected immediately after the experiment and cells were fixed with 4% polyoxymethylene for 20 min and incubated with diluted Fluo-8 AM for 30 min. After incubation, use PBS to wash the cells for three times and then use a fluorescent microscope to observe. Finally, the ratio of calcium area to the nucleus area was calculated by ImageJ.

### Transwell invasion assay

Cell invasion ability was determined by the transwell invasion assay. The transwell chambers (8.0-μm pore size; Millipore, Billerica, MA, USA) were coated with the Matrigel (diluted at 1:10; Corning Inc., USA). The cells were seeded at a density of 30,000 cells per chamber into the upper Matrigel-coated chamber and 500 μl the RPMI 1640 medium containing 10% serum was added to the bottom chamber. Then cells were exposed to the different experimental conditions (control, MB, US and US+MB). 48 h after the experiment, the cells in the upper chamber were removed with cotton swabs, and the cells in the bottom chamber were fixed in 4% polyoxymethylene for 30 min and then stained with 0.1% crystal violet (Beyotime, China) for 30 min. The ratio of violet area was calculated by ImageJ and the ratio was considered as the invasion ability.

### Statistical analysis

The experiments were performed at least three times. Quantitative data are presented as means ± standard deviation. Group means were compared using one-way analysis of variance (ANOVA) under the assumptions of normality and equal variance. Multiple comparisons were performed with Tukey correction. *P*-values <0.05 were considered statistically significant. Statistical analysis was performed using GraphPad Prism 8 (GraphPad Software Inc., CA, USA).

## Results and Discussion

### Cavitation decreases the viability and promotes the apoptosis of AsPC-1 cells

This ultrasound treatment equipment consisted of a signal generator and amplifier, a 360-kHz transducer, and a water bath (Fig. 1a). The CCK8 results showed that US and US+MB could decrease the cell viability. US irradiation alone had limited effect on cell viability (92.1±1.4%) while US+MB could effectively kill cells (70.4±1.4%), revealing that microbubbles could enhance the effect of ultrasound in decreasing cell viability (Fig. 1b).

**Fig. 1.**
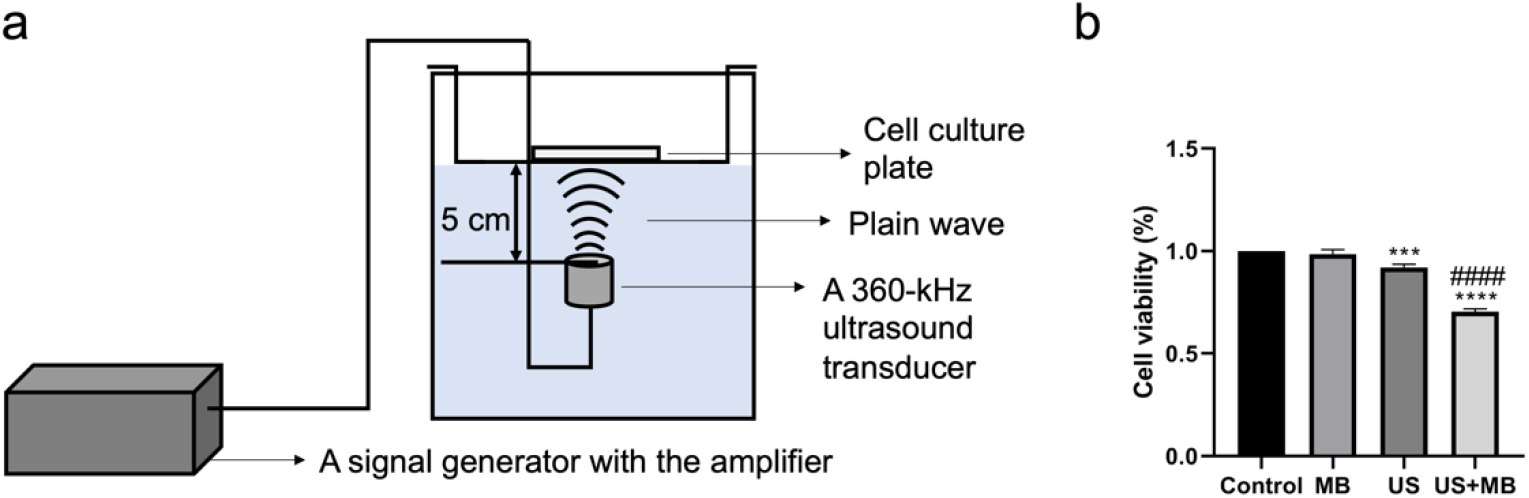
Cavitation decreased cell viability. **a** Ultrasound treatment equipment consisted of a signal generator and amplifier, a 360-kHz transducer, and a water bath. **b** Cell viability in four groups (control, MB, US, US+MB) was measured by CCK8 assay 24 h after the treatment. The optical density (OD) values are normalized by the OD value in the control group. Error bars indicated standard deviations. \*\*\*\**p*<0.0001, \*\*\**p*<0.001 vs. control; ####*p*<0.0001 vs. US; N=3. MB, microbubble only; US, ultrasound only; US+MB, ultrasound combined with microbubbles.

To investigate the mechanism underlying the decreased cell viability, we firstly examined the expression levels of the apoptosis-related proteins. Western blot analysis confirmed that cleaved Caspase-3 increased when the cells were exposed to US and US+MB and cells exposed to US+MB expressed the highest level of cleaved Caspase-3. Also, cells that treated with US+MB presented a significantly higher Bax/Bcl-2 ratio, which indicated a higher apoptosis potential of cells in US+MB group (*P*<0.05) (Fig. 2a). To explore the upstream signals of Caspase-3, we measured the expression levels of intracellular calcium and cleaved Caspase-8. Calcium imaging showed that cells exposed to US and US+MB had an increased level of intracellular calcium and cells exposed to US+MB had the highest level of intracellular calcium (Fig. 2b). However, it was observed that Caspase-8 activation was suppressed in the US+MB group and cells in the US+MB group had the lowest level of cleaved Caspase-8 (Fig. 2a). We then used flow cytometry to quantify cells at different stages of apoptosis. The analysis confirmed that more cells were in US (22.5±3.7%) and US+MB (52.3±7.8%) groups than among the cells in the control (7.8±0.4%) and MB (7.7±0.2%) groups, indicating that microbubbles enhanced ultrasound-induced cell apoptosis (Fig. 2c).

**Fig. 2.**
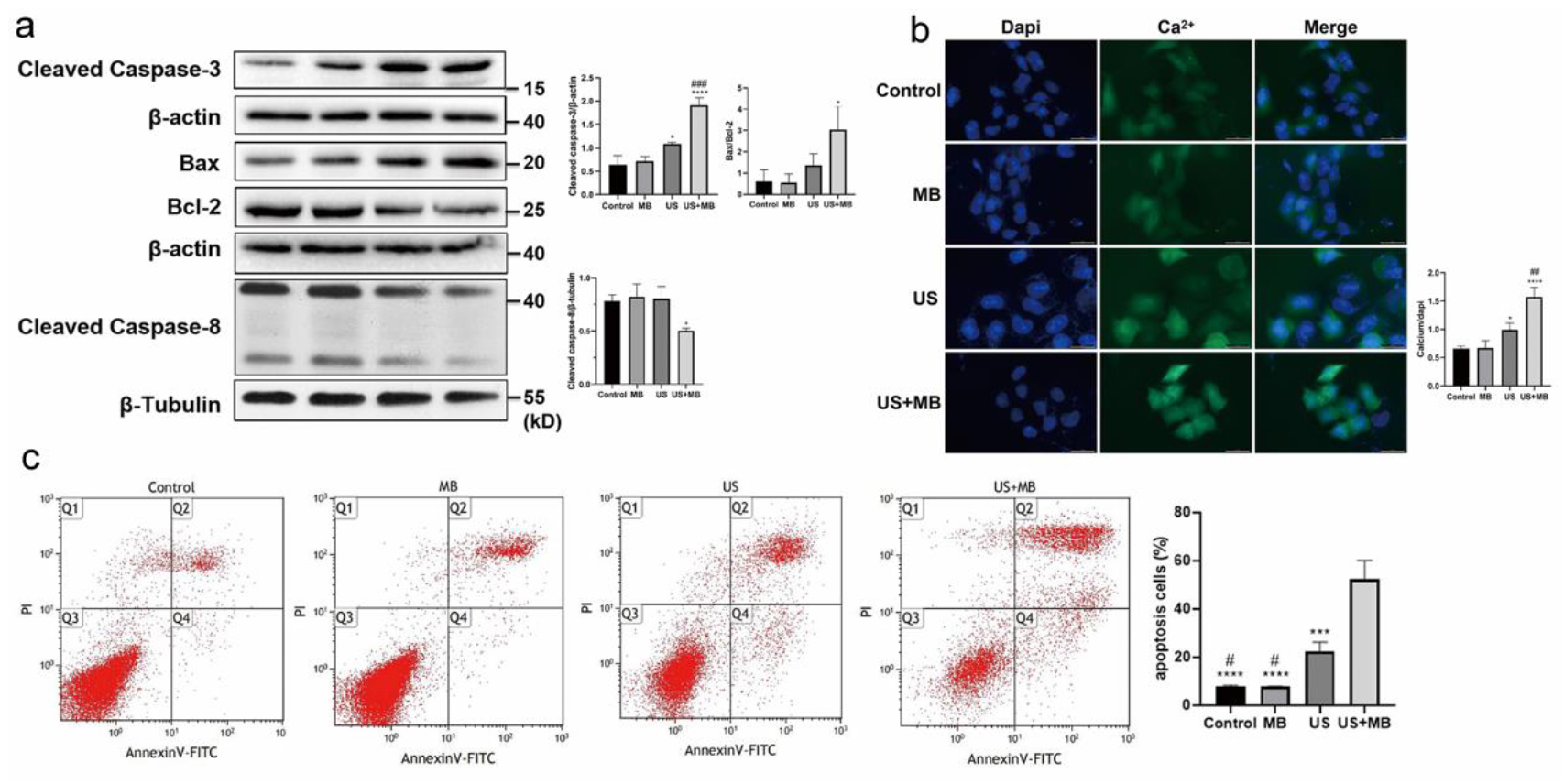
Cavitation regulated apoptosis associated proteins and promoted cell apoptosis. **a** Western blot and quantified protein levels of cleaved Caspase-3, cleaved Caspase-8, Bcl-2 and Bax in four groups (control, MB, US, US+MB) 24 h after the treatment. **b** Representative fluorescence photomicrographs of intracellular calcium load (green) analysis measured by Fluo-8 AM immediately after the treatment in four groups (control, MB, US, US+MB). Scale bars = 50 μm. **c** The ratio of cells in different apoptosis stages was obtained by flow cytometry 24 h after the treatment in four groups (control, MB, US, US+MB). Error bars indicate standard deviations. Error bars indicated standard deviations. \*\*\*\**p*<0.0001, \*\*\**p*<0.001, \*\**p*<0.01, \**p*<0.05 vs. control; ###*p*<0.001, #*p*<0.05 vs. US; N=3. MB, microbubble only; US, ultrasound only; US+MB, ultrasound combined with microbubbles. Bcl-2, B-cell lymphoma-2; Bax, Bcl-2-associated X protein.

SEM was performed to observe the morphological change on the cell membrane. Based on the results, the cell membrane was mechanically damaged and formed cell debris after exposing to US+MB. However, it was not observed that cell membrane was disrupted in the other three groups (control group, MB group and US group), indicating that shock waves mediated by microbubbles could cause severe disruption on the cell membrane (Fig. 3).

**Fig. 3.**
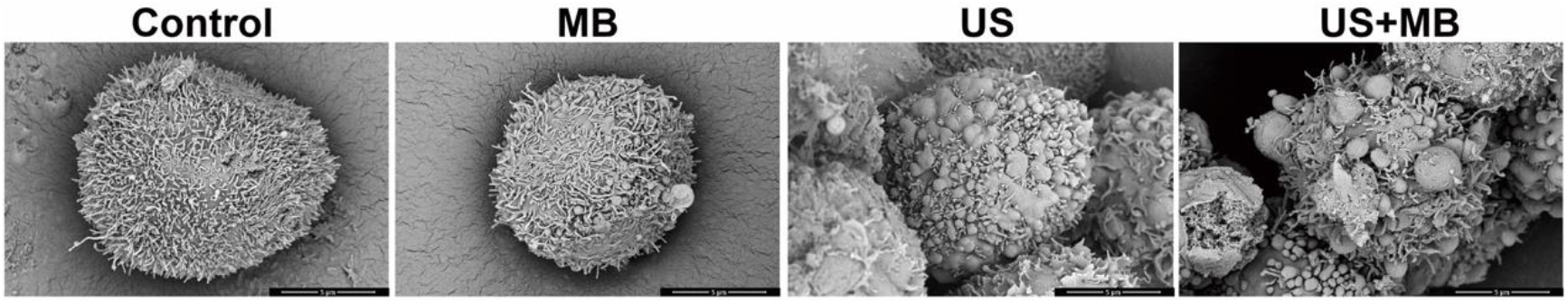
Cavitation induced mechanical disruption on cell membrane. Scanning electron microscope was used to observe the morphological change of cell membrane in four groups (control, MB, US, US+MB) immediately after the conduction (magnification, ×10000). Membrane disruption was observed in US+MB group. Scale bars = 5 μm. MB, microbubble only; US, ultrasound only; US+MB, ultrasound combined with microbubbles.

### Cavitation suppresses abilities of proliferation and colony formation of AsPC-1 cells

To measure the cell proliferation ability in four groups, we tested the Ki67 expression level using the western blot. The analysis showed that cells in the US+MB group had a lower expression of Ki67 than cells in the other three groups (*P*<0.001) (Fig. 4a). Results of the immunofluorescence staining of Ki67 were consistent with those of the western blot analysis, showing that cells in the US+MB group had the least expression of Ki67 localizing in nucleus (Fig. 4b). We also performed the colony formation assay and found that cells exposed to US+MB formed the least colony area after the 14-day culture (*P*<0.0001) (Fig. 4c).

**Fig. 4.**
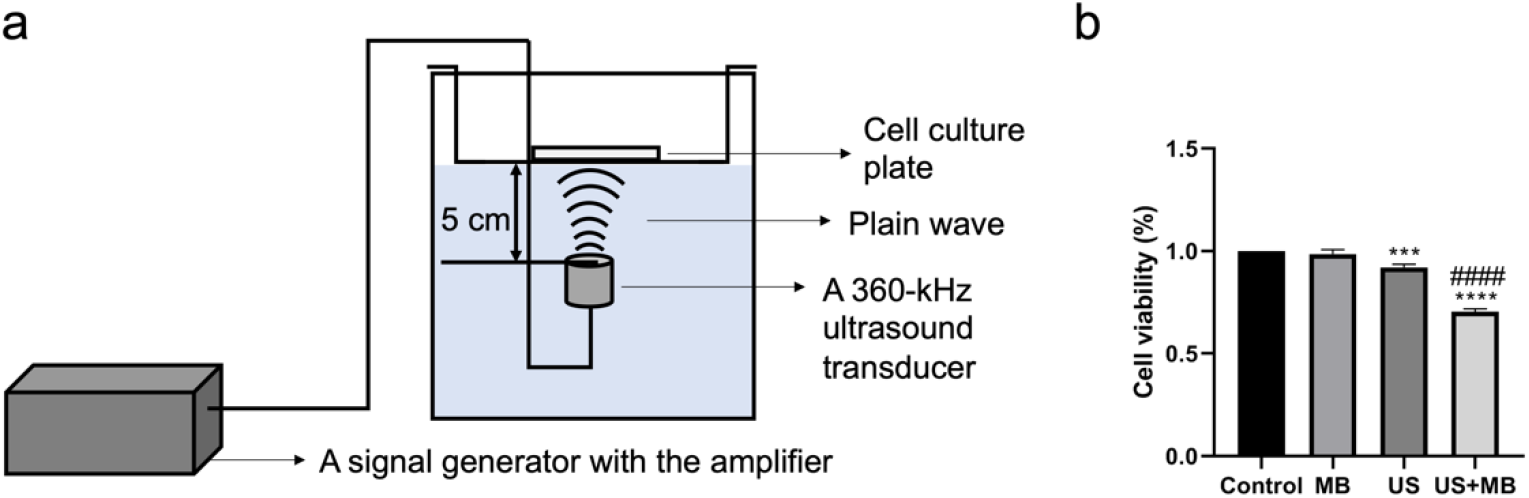
Cavitation suppressed cell proliferation ability. **a** Western blot of Ki67 24 h after the treatment in four groups (control, MB, US, US+MB). **b** Representative fluorescence photomicrographs of Ki67 (red) and nucleus (blue) after 24 h of treatment in four groups (control, MB, US, US+MB). Scale bars = 20 μm. **c** AsPC-1 cell colony formations in four groups (control, MB, US, US+MB). Cells were cultured for 14 days after exposing to different experimental conditions. Error bars indicated standard deviations. \*\*\*\**p*<0.0001, \*\*\**p*<0.001, \*\**p*<0.01 vs. control; #### *p*<0.0001, ###*p*<0.001, # *p*<0.05 vs. US; N=3. MB, microbubble only; US, ultrasound only; US+MB, ultrasound combined with microbubbles.

### Cavitation weakens invasion ability of AsPC-1 cells and down-regulates EMT associated proteins

To investigate the invasion ability of cells under the different experimental conditions, we used the transwell invasion assay and observed the cells that migrated through the polycarbonate membrane. After 48 h, cells exposed to US+MB had the least invaded cells compared with the other groups (Fig. 5).

**Fig. 5.**
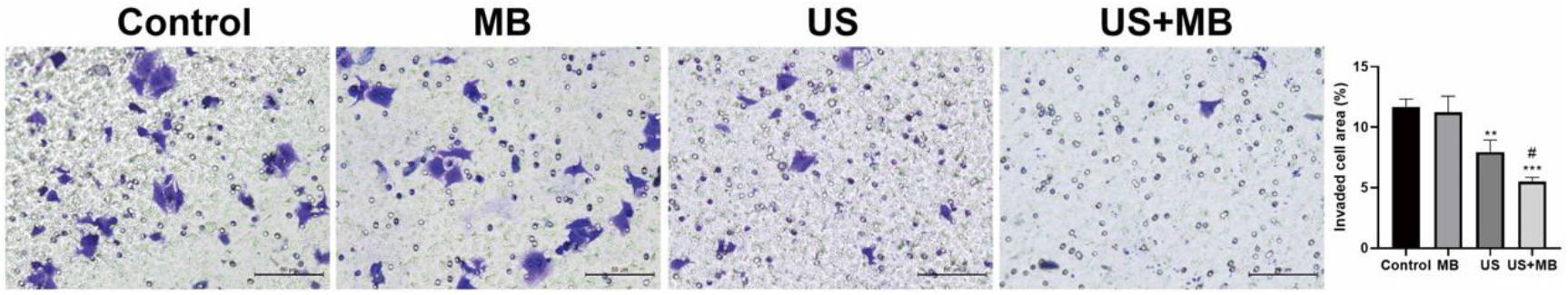
Cavitation restrained cell invasion ability. Transwell invasion assay in four groups (control, MB, US, US+MB). Cells were seeded with density of 30,000 per chamber into the upper chamber after exposing to different experimental conditions. After cells were cultured for 48 h, US+MB group showed decreased invaded cells than the other three groups. Error bars indicated standard deviations. Scale bars = 50 μm. \*\*\**p*<0.001, \*\**p*<0.01 vs. control; #*p*<0.05 vs. US; N=3. MB, microbubble only; US, ultrasound only; US+MB, ultrasound combined with microbubbles.

EMT is known to play an important role in cancer cell invasion. We examined the EMT associated proteins to determine whether EMT was suppressed by cavitation. The immunofluorescence technique was used to observe the EMT related protein expression, E-cadherin and Vimentin. We observed an increased expression level of E-cadherin and a decreased expression level of vimentin in the US+MB group comparing with the other three groups, which meant a repressed development of EMT (Fig. 6a, b). We then used western blot analysis to confirm the expression level of E-cadherin and found the highest level of E-cadherin in US+MB group (Fig. 6c), which was consistent with the immunofluorescence result. We also tested the expression level of HIF-1α and found that cells exposed to US+MB had the lowest HIF-1α expression level (Fig. 6c). These findings revealed that cavitation restrained the development of EMT and hypoxia to decrease the invasion ability in AsPC-1 cells.

**Fig. 6.**
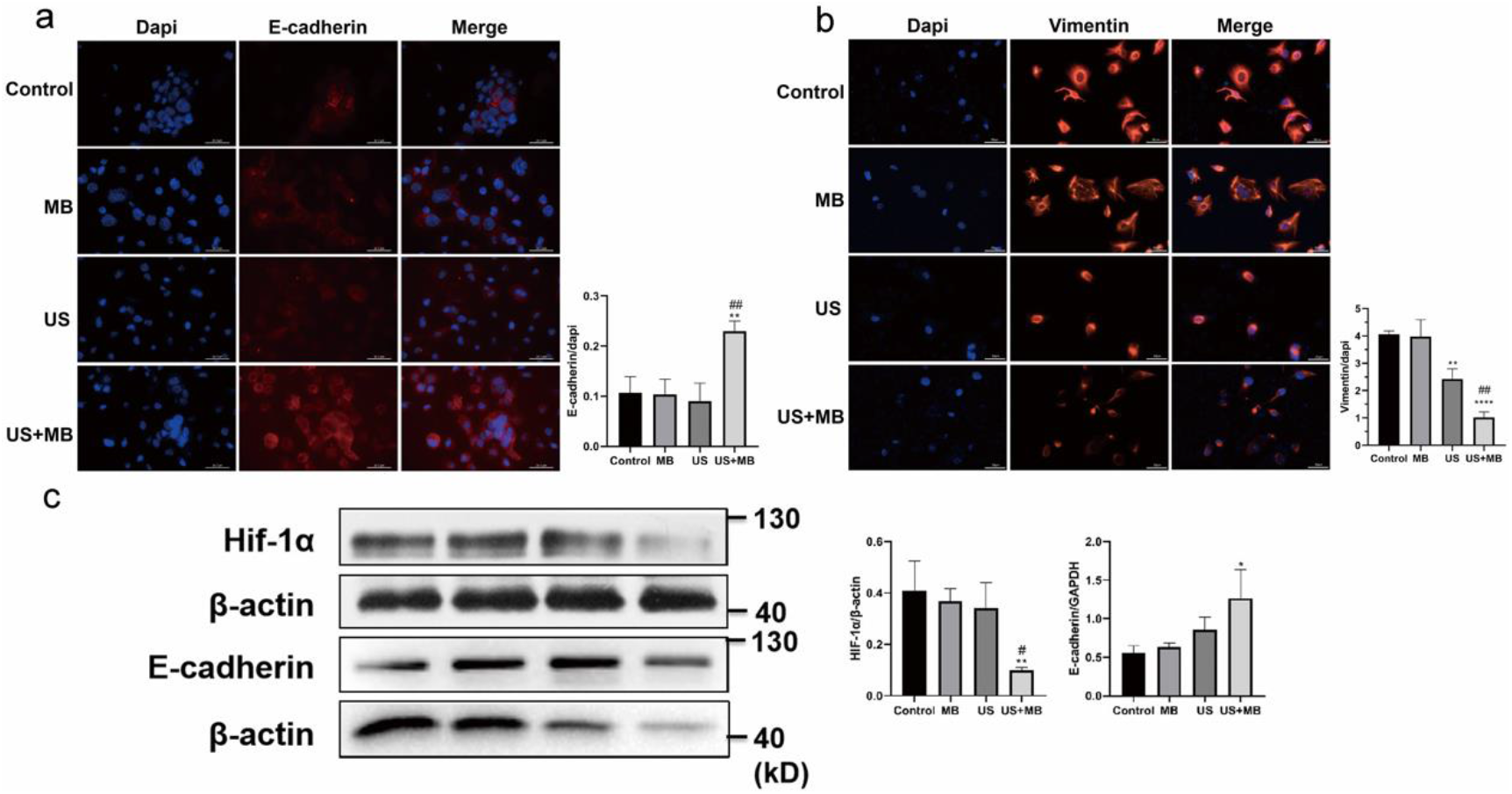
Cavitation suppressed the development of EMT and hypoxia. **a** Representative fluorescence photomicrographs of E-cadherin (red) and nucleus (blue) 24 h after the treatment in four groups (control, MB, US, US+MB). Scale bars = 50 μm. **b** Representative fluorescence photomicrographs of Vimentin (red) and nucleus (blue) 24 h after the treatment in four groups (control, MB, US, US+MB). Scale bars = 50 μm. **c** Western blot was used to measure expression level of HIF-1α and E-cadherin after 24 h of treatment in four groups (control, MB, US, US+MB). Error bars indicated standard deviations. \*\**p*<0.01, \**p*<0.05 vs. control; ## *p*<0.01, # *p*<0.05 vs. US; N=3. MB, microbubble only; US, ultrasound only; US+MB, ultrasound combined with microbubbles. HIF-1α, hypoxia-inducible factor-1α.

### Cavitation reduces invadopodia formation in AsPC-1 cells

Finally, we investigated the invadopodia formation, which are dynamic actin-rich protrusions that degrade the extracellular matrix and are involved in cancer cell metastasis (20). As shown by arrows, fewer invadopodia were observed in the US+MB group (Fig. 7).

**Fig. 7.**
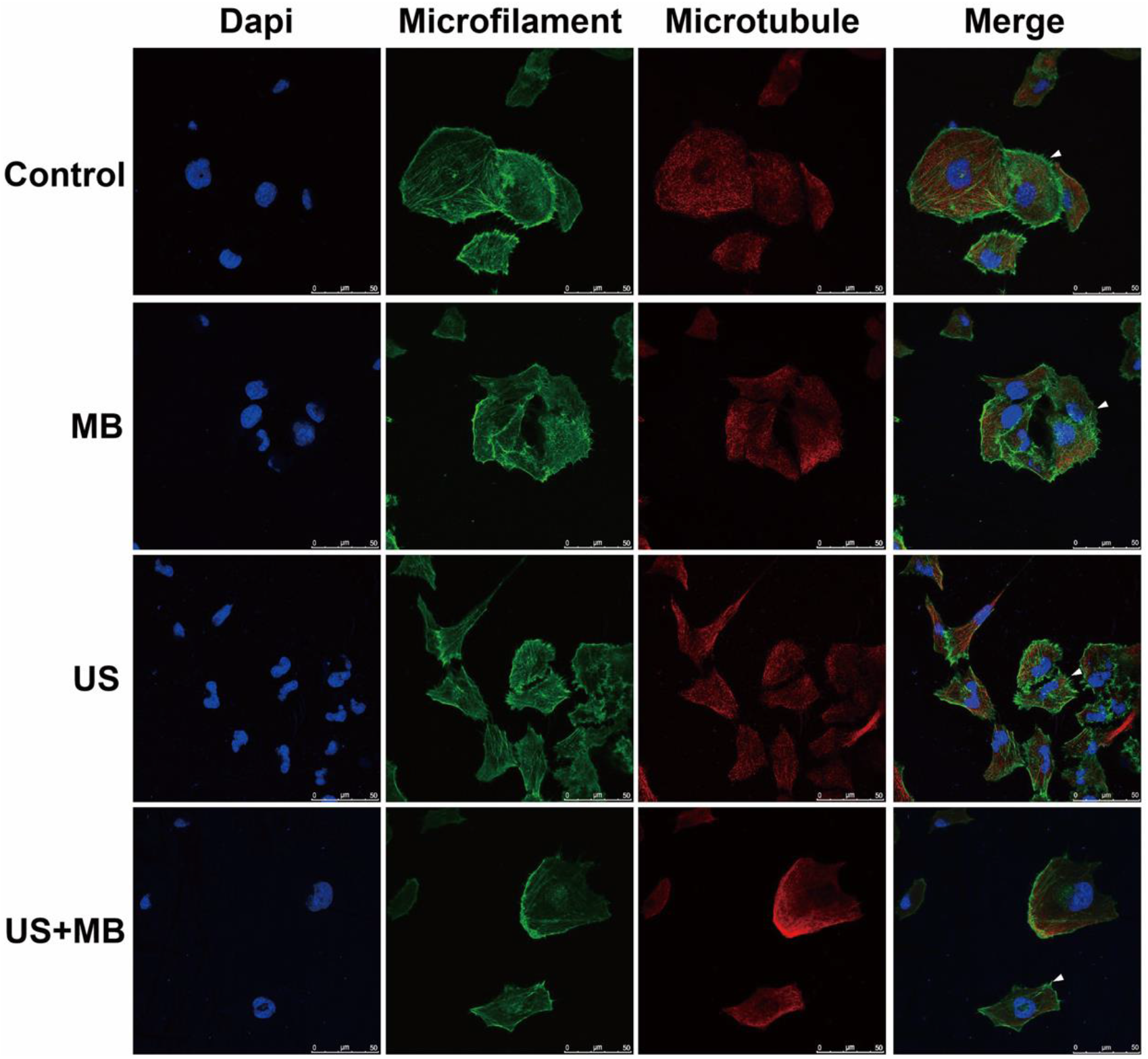
Cavitation reduced invadopodia. Representative fluorescence photomicrographs of microfilament (red), microtubule (green) and nucleus (blue) revealed the morphology of cytoskeleton and invadopodia (indicated by arrow) after 24 h of treatment in four groups (control, MB, US, US+MB). Cells in US+MB group had fewer invadopodia. Scale bars = 50 μm. MB, microbubble only; US, ultrasound only; US+MB, ultrasound combined with microbubbles.

Acoustic cavitation can release the shock wave with the existence of microbubbles. The shock wave can attenuate cell viability through cell apoptosis and necrosis (21, 22). In this research, we ensured that both US and US+MB could decrease cell viability but US had limited effect on cell viability. SEM results confirmed the mechanical disruption on cell membrane which was caused by cavitation. Western blot analysis of cleaved Caspase-3 indicated that Caspase-3 mediated apoptosis pathway was activated when cells were conducted with cavitation. When exploring the upstream signals of Caspase-3, we found that cleaved Caspase-8 was decreased after cells were treated with US+MB. Caspase-8 is at the crossroads of necroptosis pathway and death receptor-mediated apoptosis pathway. When the death receptor-mediated pathway is activated, caspase-8 can activate caspase-3 directly or cleave the protein Bid, leading to activation of its downstream effector proteins Bax and Bak. However, active Caspase-8 is inhibited when necroptosis occurs (23). Another upstream signal of Caspase-3 is the intracellular calcium. In this research, we observed that cells both in US and US+MB groups had increased levels of intracellular calcium, which was consistent with the results of previous researches (15, 16). Therefore, the decreased level of cleaved Caspase-8 and the increased intracellular calcium in US+MB group indicated that cavitation-induced Cacpase-3 active was mainly due to the increased intracellular calcium. In addition, the decreased level of cleaved Caspase-8 suggested that the necroptosis pathway was enhanced by cavitation.

For cell proliferation, we used Ki67 and colony formation to evaluate cell proliferation ability. Ki67 is a protein which has a high expression level during the cell cycle and reduces when cell leaves the cell cycle (24, 25). 24 h after the treatment, we confirmed that US+MB down-regulated Ki67, which meant that more cells leaved cell cycle and entered apoptosis or necroptosis program. Colony formation reflects the characters of proliferation ability and population dependence. After a 2-week culture, we found that US+MB induced a decreased colony formation ability in AsPC-1. These finds revealed that US+MB could lower down the proliferation ability.

For cell invasion, cells that migrated through the polycarbonate membrane were determined not only by the ability of degrading the Matrigel but also by the number of cells. Therefore, the transwell invasion result in US+MB group might be caused by the collapse of cell population. This result pointed out that effective killing of cancer cells was a strategy to prevent cancer cell invasion. According to previous reports, HIF-1α is a major factor that reflects the degree of hypoxia in cells or tissues and promotes the occurrence of EMT (26, 27). In this research, we confirmed that US+MB could reduce the HIF-1α expression and relieve hypoxia in the microenvironment. The immunofluorescence imaging and western blot also provided the expression of EMT associated proteins in cellular level illustrating that cavitation could restrain EMT development. These findings suggested that cavitation could effectively reduce cancer cell invasion ability. During the process, hypoxia served a bridge which connected cavitation-induced apoptosis with decreased invasion ability in cells.

In the present study, we confirmed that the increased intracellular calcium contributed to cavitation-induced pancreatic cancer cell apoptosis and the decreased activation of Caspase-8 was supposed to related to the enhancement of necroptosis, revealing that multiple mechanisms participated in cavitation-induced cell death. We also observed that the down-regulated invasion ability of pancreatic cancer cell with a lower level of HIF-1α and a restrained EMT development after the conduction of ultrasound combined with microbubbles. But the limitation of this study is that these results are only tested in *Vitro*. Besides, role of necroptosis in cavitation-induced cell death requires further research based on the decreased activation of caspase-8 in this study.

## Conclusion

In conclusion, this research clarifies that microbubble-mediated cavitation promotes the apoptosis in AsPC-1 cells through the increased intracellular calcium and suppresses the invasion ability of AsPC-1 cells through down-regulated HIF-1α and restrained EMT development. We also find that cavitation-induced cell death is a process which multiple mechanisms participated in. Based on these findings, microbubble-mediated cavitation is a considerable method for cancer treatment.

## Author contributions

Pintong Huang designed this study. Jing Cao and Feng Diao performed the experiments. Jing Cao, Hang Zhou, Fuqiang Qiu, Jifan Chen acquired the data. Jing Cao and Hang Zhou analyzed and interpreted the data. Jing Cao wrote the main manuscript text. All authors reviewed the manuscript.

## Acknowledgements

The study was supported by the National Natural Science Foundation of China (Grant No. 81527803, 81420108018), National Key R&D Program of China (No.2018YFC0115900), and Zhejiang Science and Technology Project (No. 2019C03077). There are no competing financial interests in relation to this research.

